# Phloem-mediated spreading of SIGS-derived non-coding RNAs in *Hordeum vulgare*

**DOI:** 10.1101/2019.12.30.891002

**Authors:** D Biedenkopf, T Will, T Knauer, L Jelonek, ACU Furch, T Busche, A Koch

## Abstract

Small (s)RNA molecules are crucial factors in the communication between hosts and their interacting pathogens/pests that can modulate both host defense and microbial virulence/pathogenicity known as cross-kingdom RNA interference (ckRNAi). Consistent with this, sRNAs and their double-stranded (ds)RNA precursors have been adopted to control plant diseases through exogenously applied RNA biopesticides, known as spray-induced gene silencing (SIGS). While RNA spray proved to be effective, the mechanisms underlying the transfer and uptake of SIGS-associated RNAs are inadequately understood. Moreover, the use of the SIGS-technology as a biopesticide will require the systemic spreading of dsRNA/siRNA signals. Our results strongly support the notion of phloem-mediated long-distance movement of SIGS-associated dsRNA and/or siRNA. These findings are significant contributions to our mechanistic understanding of RNA spray technology, as our previous data indicate that SIGS requires the processing of dsRNAs by the fungal RNAi machinery. In summary, our findings support the model that SIGS involves: (i) uptake of sprayed dsRNA by the plant (via stomata); (ii) transfer of apoplastic dsRNAs into the symplast (DCL processing into siRNAs); (iii) systemic translocation of siRNA or unprocessed dsRNA via the vascular system (phloem/xylem); (iv) uptake of apoplastic dsRNA or symplastic dsRNA/siRNA depending on the lifestyle/feeding behavior of the pathogen/pest.

## Introduction

The vascular network of higher plants is composed of the phloem and xylem that pervades the whole organism from root to shoot and distributes nutrients and water (van Bel et al. 2011, Lucas et al. 2013). Sieve elements (SEs), companion cells (CCs) and phloem parenchyma cells (PPCs) are the three phloem elements involved in long-distance transport of photoassimilates in angiosperms (van Bel 1996, Hafke et al. 2005). A high density of pore-plasmodesma units (PPUs) and ER coupling between SE and CC underline an intimate symplasmic connection (Kempers et al. 1998, Martens et al. 2006). The cross-walls between the SE-modules become transformed into sieve plates, perforated by plasmodesmata (PD) modified into sieve pores, mediating long-distance transport of signalling molecules that play a pivotal role for the regulation of several developmental processes (Lough and Lucas 2006). This non-cell-autonomous control involves the transfer of informational molecules such as proteins, mRNA and small RNAs (Ham and Lucas 2017). Since the first detection of unspecified nucleic acids in the phloem sap in the late 1990s (Sasaki et al. 1998, Ruiz-Medrano et al. 1999), several studies demonstrated the systemic translocation of mRNA mediating non-cell autonomous control of plant development, defense and nutrient allocation via the phloem (Lough and Lucas 2006, Ham and Lucas 2017). However, decades ago scientists revealed RNA as the agent for systemic acquired gene silencing. They have shown delivery of RNA-based signals via the phloem pathway that affect gene expression at the whole-plant level by sequence-specific degradation of targeted mRNA (Jorgensen 1995, Palauqui et al. 1996, 1997). These scientists were the first who linked systemic RNA signaling with a process known as RNA silencing.

RNA silencing (also known as RNA interference, RNAi) is a conserved and integral aspect of gene regulation mediated by small RNAs (sRNAs) that direct gene silencing on the level of transcription but also post-transcriptionally. At the transcriptional level, gene expression is inhibited via RNA-directed DNA methylation (RdDM) while at the post-transcriptional level (PTGS) direct mRNA interference causes inhibition of translation. Originally, RNA silencing is associated with protection against viral infection, control of epigenetic modifications, regulation of genome stability, curbing of transposon movement and regulation of heterochromatin formation (Castel and Martienssen 2013; Koch et al. 2017). Besides its natural function, RNA silencing has emerged as a powerful genetic tool for scientific research over the past several years. It has been utilized not only in fundamental research for the assessment of gene function, but also in various fields of applied research, such as agriculture. In plants, RNA silencing strategies have the potential to protect host plants against predation or infection by pathogens and pests mediated by lethal RNA silencing signals generated *in planta* (for review see: Koch and Kogel 2014; Yin and Hulbert 2015; Zhang et al., 2017; Qi et al., 2019; Liu et al. 2019). Indeed, our results showed that transgenic *Arabidopsis* and barley (*Hordeum vulgare*) plants, expressing a 791 nucleotide (nt) dsRNA (CYP3RNA) targeting all three copies of the *CYP51* gene (*FgCYP51A*, *FgCYP51B*, *FgCYP51C*) in *Fusarium graminearum* (*Fg*), inhibited fungal infection via a process designated as host-induced gene silencing (HIGS) (Nowara et al. 2010, Koch et al. 2013). Moreover, we demonstrated that HIGS-mediated targeting of the structural sheath protein (Shp) mandatory for aphid feeding, produced significantly lower levels of Shp mRNA compared to aphids feeding on wild-type (wt) plants (Abdellatef et al. 2015).

In addition to the generation of RNA silencing signals *in planta*, plants can be protected from pathogens and pests by exogenously applied RNA biopesticides (known as spray-induced gene silencing, SIGS) (for review see: Mitter et al. 2017; Cai et al. 2018; Dubrovina and Kiselev 2019; Gaffar and Koch 2019; Dalakouras et al. 2020). Over the last decade, our research (Koch et al., 2013; 2016; 2018; 2019; Höfle et al., 2019, Werner et al., 2019) and the work of others have revealed the enormous potential of RNA-silencing strategies as a potential alternative to conventional pesticides for plant protection, regardless of how target-specific inhibitory RNAs are applied (i.e. endogenously or exogenously).

Despite the promising potential of RNAi-based disease control and its benefits for agronomy and the ecosystem, the mechanisms underlying those technologies are virtually unresolved. Understanding the uptake and translocation processes of non-coding RNAs is critical for its successful future application in the field. Application non-coding RNAs as biopesticides will require knowledge on the paths used by dsRNA/siRNA as signal. Previously, we have shown that fluorescent dsRNA is detected in the vascular tissue of barley after spraying the leaves with the 791 nt long CYP3-dsRNA_A488_, directed against fungal *CYP51* genes using a detached leaf assay that enabled us to assess fungal growth in distal (semi-systemic, non-sprayed) leaf segments (Koch et al. 2016).

In the present study, we demonstrated that aphids accumulate fluorescent dsRNA, when they feed on distal parts of barley leaves that were sprayed with a Shp-dsRNA. Our study claimed to further prove that sprayed RNAs are systemically translocated via the phloem. Summarizing our results we (i) found spray-induced gene silencing of an aphid mRNA target measured by qRT-PCR, (ii) visualized a fluorescent-labelled dsRNA in the phloem sap of barley using stylectomy (iii) profiled SIGS-derived siRNA in the phloem sap of barley by RNA-Seq analysis (iv) detected SIGS-associated RNAs in barley roots using RT-PCR.

## Material and Methods

### Maintenance of plants and aphids

The spring barley (*Hordeum vulgare*) cultivar (cv.) Golden Promise (GP) was grown in a climate chamber under 16 h light photoperiod (240 μmol m^−2^ s^−1^ photon flux density) at 18°C/14°C (day/night) and 65% relative humidity. Grain aphids (*Sitobion avenae*) were reared on three-week-old barley plants in a climate chamber under the same conditions. To obtain synchronized insects, reproductive mature aphids were placed in clip cages (one aphid per cage) on GP plants for 24 h. The adults were then removed, and the offspring were used for experiments as previously described (Gaupels et al., 2008; Schmitz et al., 2012).

### dsRNA synthesis

For spray experiments, the clone p7i-Ubi-Shp-RNAi that includes a partial sequence of the 3,621 bp *Shp*-cDNA (XM_001943863, ACYPI009881) of the pea aphid *Acyrthosiphon pisum* (*Ap*) was used as template for the synthesis of 491 nt long Shp-dsRNA (Abdellatef et al., 2015). dsRNA was generated using MEGAscript RNAi Kit (Invitrogen) following MEGAscript^®^ protocols. Primer pairs T7_F and T7_R with T7 promoter sequence at the 5` end of both forward and reverse primers were designed for amplification of dsRNA (S1 Table).

### Spray application of dsRNA

Second leaves of 2-3-wk-old barley cv. GP were detached and transferred to square petri plates containing 1% water-agar. The dsRNA was diluted in 500 μl water to a final concentration of 20 ng μl^−1^. As a control, TRIS/EDTA (TE) buffer was used in a concentration corresponding to final concentration of the dsRNA sample. A typical RNA concentration after elution (see MEGAscript^®^ protocols) was 500 ng μl^−1^ with a final TE buffer concentration of 400 μM Tris-HCL and 40 μM EDTA. Spraying of leaves was carried out using a spray flask as described (Koch et al., 2016). Each plate containing 10 detached leaves were sprayed in a semi-systemic setup where lower leaf segments were covered with a plastic tray as described (Koch et al., 2016), with either Shp-dsRNA or TE buffer by giving 3-4 puffs, and subsequently kept at room temperature (RT). Forty-eight hours after spraying, aphids were placed on the non-sprayed part of each leaf using clip cages.

For the RNA translocation assays barley seedling were grown in petri dishes for one week. The seedlings were then placed into moist filter paper rolls and grown for 4 days before spraying the first leaf with 20 ng μl^−1^ of CYP3RNA or GFP-dsRNA (control). To measure the amount of sprayed dsRNA in different tissues (first leaf, second leaf, shoot and root), samples were taken 24h, 48h and 72h after spray treatment of the first leaf. To analyse target gene silencing plants were inoculated with *Fg* after spray treatment and samples were taken five days after infection.

### Stylectomy and aphid sampling

At 24 h after spray application of dsRNA, 20-30 mainly adult aphids (*Sitobion avenae*) were placed onto the upper surface of each leaf and allowed to feed for 24 h. Following successful cauterization of the mouthparts with a microcautery device (CF-50, Syntech, Fisher and Frame 1984), each aphid stylet exuding phloem sap was marked with two small dots on the leaf surface and kept moist with a drop of DEPC water to prevent early occlusion. After cutting all aphid stylets, the DEPC water droplets were removed with a paper towel and the petri dish was subsequently flooded with silicone oil (M 200, Roth) to prevent evaporation of the sieve tube exudates. Due to the observation of sporadic bacterial and fungal contamination from the leaf surface, a mixture of antimycotic (25 nl Nystatin 5mM), antibiotic (25 nl Tetracycline 5 mM), and 50nl RNAse Inhibtor (Invitrogen) (with traces of Bromophenol Blue for optical verification) was injected into each sieve tube sap sample at the beginning of the exudation phase to prevent degradation of RNA. By 24 h later, the sieve tube sap was collected using a microcapillary connected to a small syringe via a silicone tube with a side valve. Depending on the exudation time of the severed stylets, the sample amount reached up to 2 μl per stylet over 24 h. The samples of each treatment were pooled and stored at −80°C.

### Aphid transcript analysis

To assess silencing of the *Shp* gene, mRNA expression analysis was performed using quantitative real-time PCR (qRT-PCR). Freshly extracted mRNA from 10 aphids was converted into cDNA using the QuantiTect Reverse Transcription Kit (Qiagen, Hilden, Germany) and 40 ng of cDNA was used as the template for qRT-PCR in an Applied Biosystems 7500 FAST real-time PCR system. Each reaction comprised 7.5 μl SYBER Green JumpStart Taq ReadyMix (Sigma-Aldrich, Steinheim, Germany) and 0.5 pmol of the gene-specific primers Shp-RNA-qpcr-F1 and Shp-RNA-qpcr-R1 (Table S1). After initial heating to 95°C for 5 min, the target was amplified by 40 cycles at 95°C for 30 s, 52°C for 30 s and 72°C for 30 s. Ct values were determined with the 7500 Fast software supplied with the instrument. Levels of *Shp* transcripts were determined via the 2^−∆∆Ct^ method (Livak and Schmittgen 2001) by normalizing the amount of target transcript to the amount of the reference transcript 18S ribosomal RNA (GenBank APU27819).

### Small RNA library production and sequence analysis

RNA enriched for the sRNA fraction was purified from plant and fungal samples using the mirVana miRNA Isolation Kit (Life Technologies). Indexed sRNA libraries were constructed from these enriched sRNA fractions with the NEBNext Multiplex Small RNA Library Prep Set for Illumina (New England Biolabs) according to the manufacturer’s instructions. Indexed sRNA libraries were pooled and sequenced on the Illumina HiSeq and NextSeq 500 platforms and the sequences sorted into individual datasets based on the unique indices of each sRNA library. The adapters and indices were trimmed with Cutadapt (Martin 2011) version 1.16. Only reads with a length between 19 bp and 30 bp in the first pair of the paired end dataset were analysed. The reads were mapped to the *shp*-dsRNA vector sequence using bowtie2 (Langmead and Salzberg 2012) with “-very-sensitive −L 10” to identify sRNAs with a perfect match. The libraries contained 4.1 and 3.5 (control) million reads before trimming and filtering and 0.6 and 0.3 million reads after trimming and filtering.

### Confocal microscopy of fluorophore distribution

Fluorescent labeling of dsRNA was performed using the Atto 488 RNA Labeling Kit (Jena Bioscience, Jena, Germany) following the manufacturer’s instructions. Leaves were sprayed in a semi-systemic setup with the labeled dsRNA. Twenty-four h after spraying of fluorescing dsRNA, aphids were placed onto the leaves and allowed to feed overnight. Twenty-four h after infestation only stylets of the non-sprayed leaf area were cut. Phloem droplets that appeared on the stylet tips were imaged using a Leica TCS SP2 (Leica Microsystems, Wetzlar, Germany) equipped with a 75-mW argon/krypton laser (Omnichrome, Chino, CA) and a water immersion objective (HCX APO L40×0.80 W U-V-l objective).

## Results

### SIGS-mediated gene silencing in *Sitobion avenae* fed on barley leaves

Mobile cell non-autonomous inhibitory RNAs that spread gene silencing into adjacent cells and tissues have been shown to move through the vascular system (Lewsey et al., 2016, Palauqui et al., 1997, Zhang et al., 2016). Previously, we demonstrated that aphids which fed on transgenic barley expressing dsRNA derived from the *Shp* gene (Shp-dsRNA), produce significantly lower levels of *Shp* mRNA compared to aphids feeding on wild-type (wt) plants (Abdellatef et al., 2015). Based on these previous data, we modified the setup for dsRNA application in order to demonstrate a semi-systemic transfer of the dsRNA. Therefore, we tested whether locally sprayed 491 nt long Shp-dsRNA confers gene silencing in grain aphids feeding from distal, non-sprayed segments of the same barley leaves. To this end, the upper part of detached leaves (local tissue) was sprayed with 20 ng μL^−1^ Shp-dsRNA, while the lower part (distal tissue) was covered by a plastic tray to prevent direct dsRNA contact. After 48 h, we placed aphids at the distal, non-sprayed part of the leaves in clip cages and led them feed on the phloem for 24 h. Subsequently aphids were harvested and assessed for downregulation of the *Shp* target gene expression using qRT-PCR. The relative expression level of the aphid’s *Shp* gene was reduced by almost 60% compared to aphids that fed on control leaves sprayed with Tris-EDTA (TE) buffer (Fig. 1), suggesting that the transfer of inhibitory RNA from the plant phloem sap to the insect was successful.

**Figure 1:**
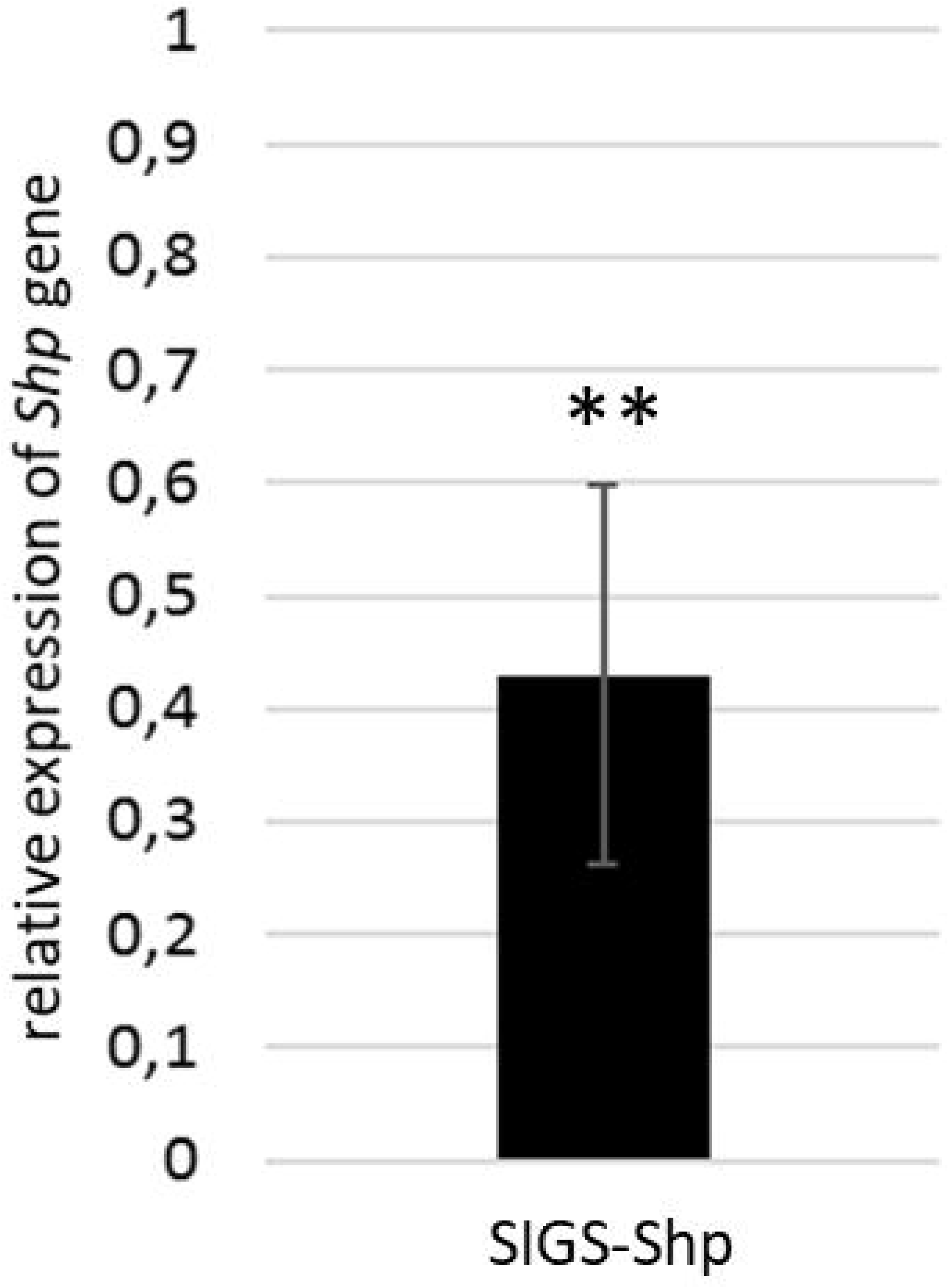
Silencing of *Shp* gene in *Sitobion avenae* that fed on dsRNA sprayed barley leaves. Gene-specific expression of *Shp* was measured by qRT-PCR and normalized to aphid 18S ribosomal RNA (GenBank APU27819) as reference gene. cDNA was generated after total RNA extraction from aphids at 5 days of inoculation after dsRNA spray. The reduction in *Shp* gene expression in the aphids that fed on Shp-dsRNA sprayed leaves compared to the mock sprayed control was statistically significant. Error bars represent SE of three independent experiments each using 5-10 adult aphids for each treatment (**p<0.01; students t-test).

### Transport of sprayed dsRNA in the phloem of barley

Encouraged by the RNAi effects on the *Shp* target gene in phloem-sucking *S. avenae*, we investigate whether the sprayed Shp-dsRNA is translocated in the phloem and/or processed by the plant’s silencing machinery. To this end, we used aphid stylectomy to gain access to pure phloem sap of barley leaves (Fig. 2 A-C). Stylectomy is commonly used to study a broad variety of physiological, mechanical and molecular properties of the plant phloem (Turgeon and Wolf 2009; Peel 1975; Thompson and van Bel 2012).

**Figure 2:**
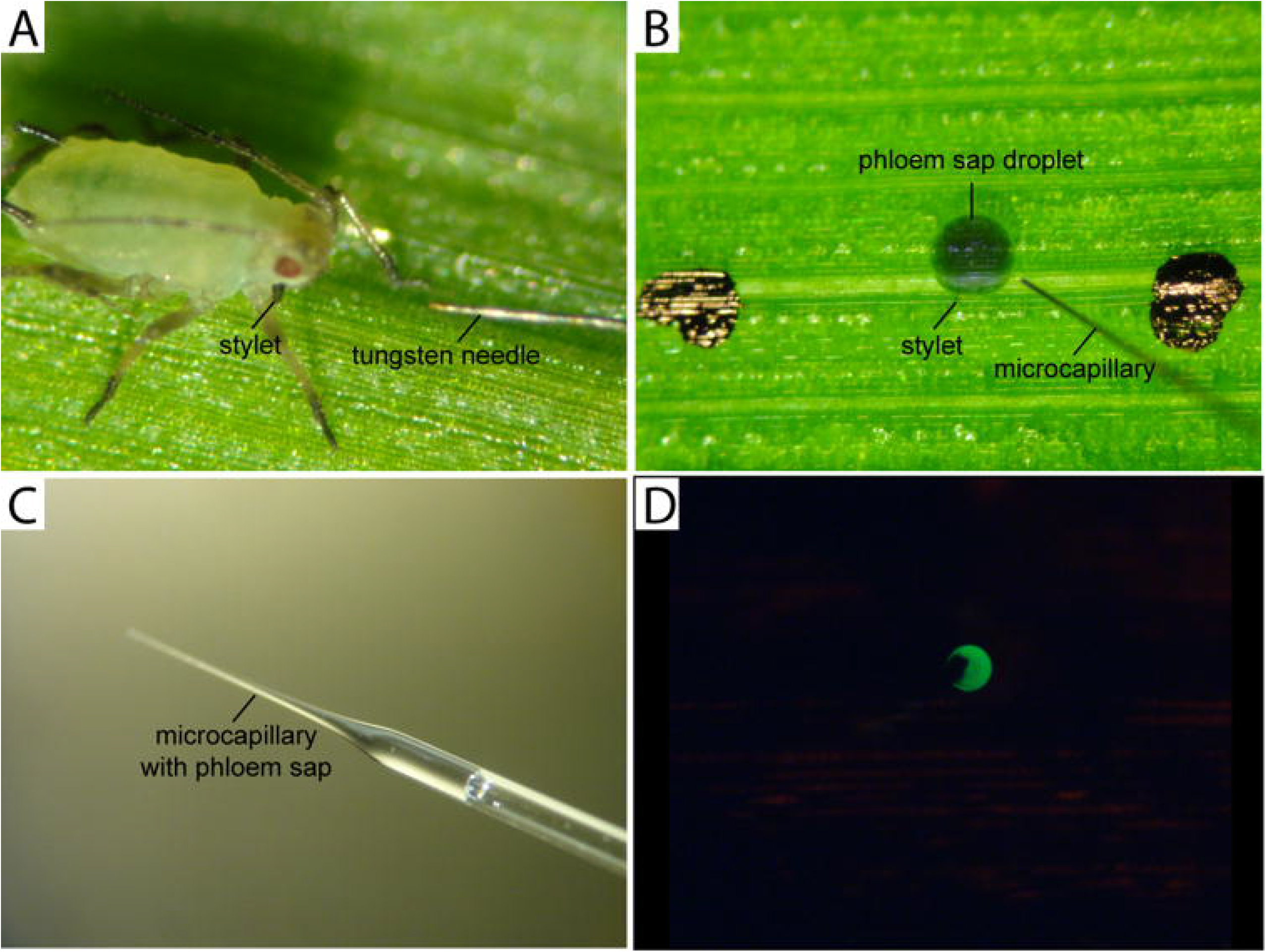
Stylectomy. (A) *Sitobion avenae* feeding on a barley leaf shortly before cutting the mouthparts (stylet) with a tungsten needle connected to a microcautery device. (B) Injection of antimycotic, antibiotic and RNAse-inhibitor using a microcapillary at the beginning of the exudation period after overlaying with silicone oil. (C) Sampling of sieve-tube exudate from a cut aphid stylet under silicone oil with a microcapillary after an exudation period of 24 hours. (D) Stylectomy of ATTO 488-labeled Shp-dsRNA_A488_ in sprayed barley leaves. Detection of Shp-dsRNA_A488_ (green) in the phloem sap droplet using stylectomy. Aphids were placed on non-sprayed leaf parts and stylectomy was performed 48 hours after spray treatment with Shp-dsRNA_A488_.

To further visualize the phloem-mediated transfer of sprayed Shp-dsRNA, it was labeled with the green fluorescent dye ATTO 488 (Shp-dsRNA_A488_) and sprayed onto barley leaves using the semi-systemic setup followed by stylectomy with *S. avenae* in the distal, non-sprayed leaf parts. Using confocal laser scanning microscopy, a green fluorescent signal was detected 24 h after feeding (48 h after spraying) and cutting at the stylet tip (Fig. 2 D). Together these data show that sprayed Shp-dsRNA is transferred via the plant’s phloem.

### RNA-Seq profiling of stylectomy samples revealed SIGS-derived siRNAs

We addressed the question whether the phloem-transferred Shp-dsRNA is stable during transport or alternatively at least partially processed into small interfering RNAs. To test this possibility, we profiled Shp*-*dsRNA-derived siRNAs in the phloem sap after isolation using stylectomy. Small RNA sequencing (sRNAseq) analysis revealed Shp-dsRNA-derived siRNA in distal (non-sprayed) leaf segments (Fig. 3). These data suggest that Shp-dsRNA-derived siRNAs also are processed by the plant’s silencing machinery and systemically transferred via the phloem.

**Figure 3:**
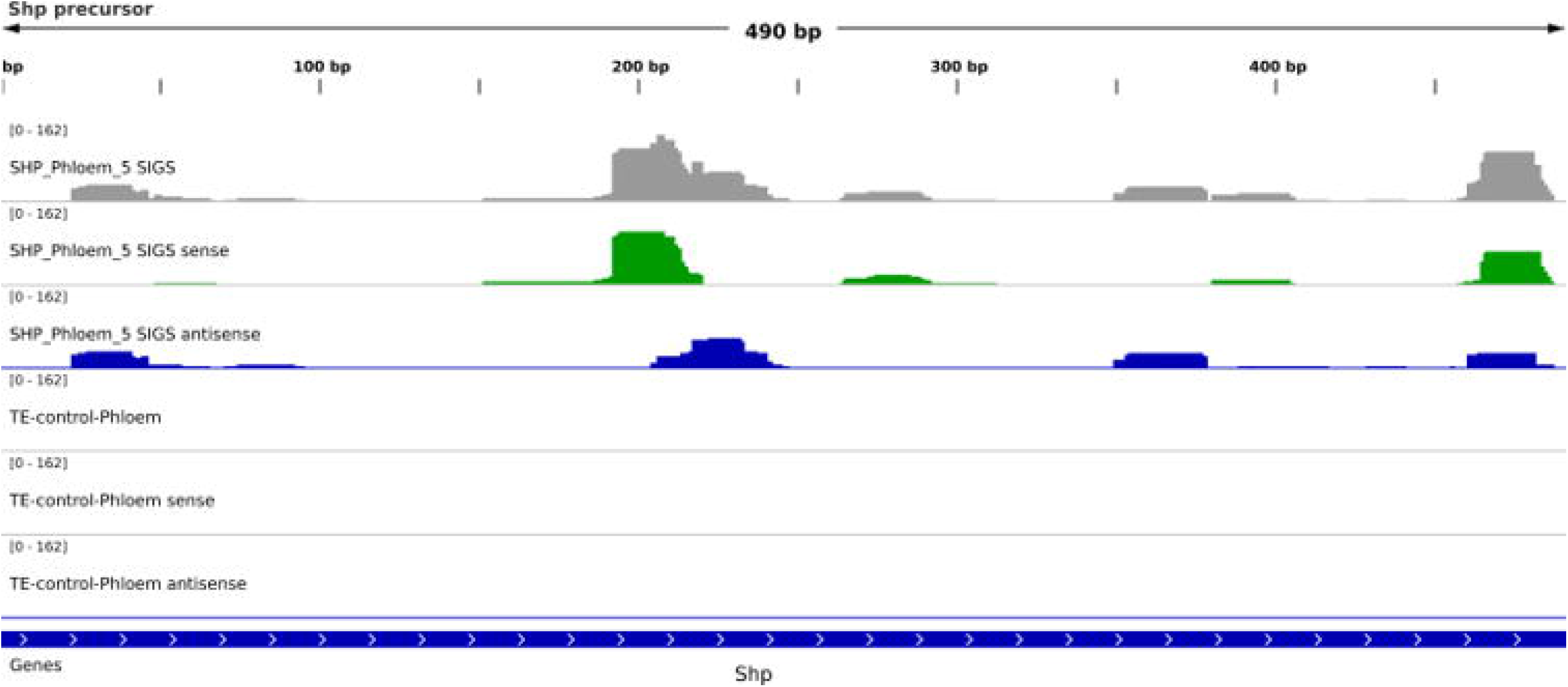
Profiling of Shp-dsRNA-derived siRNAs in the phloem sap from barley leaves. Phloem sap was sampled by stylectomy. Total RNAs were isolated using a single cell RNA purification kit (Norgen Biotek Corp.). sRNA reads between 19 and 30 nt from Shp-dsRNA-sprayed and mock-treated barley leaves were mapped to Shp-dsRNA precursor (Abdellatef et al., 2015). Read coverage varied from 0-162 as indicated. Sequencing data are gained from 5 pooled separate phloem sap isolations experiments with a total sample amount of 10μl phloem sap from sprayed leaves and 10μl phloem sap from non-sprayed leaves, respectively.

### Translocation of sprayed dsRNA from leaves to barley roots

The RNA silencing signal can travel over long distances and trigger silencing in distant plant tissues (Molnar et al., 2010, Lewsey et al., 2016). However, little is known about how SIGS-derived dsRNAs and/or siRNAs are transported in the plant. To test whether there is wider systemic spreading of RNA silencing signals via the phloem and to further determine to what extend those sprayed RNAs are translocated within the plant we analysed the spreading of the sprayed dsRNA within intact barley plants. Towards this, we measured the amount of sprayed RNA in leaves, shoots and roots at different timepoints after spray application using qRT-PCR. To assess transport of sprayed dsRNA in systemic, non-sprayed plant tissue we used two-week old barley plants that were grown in filter paper. For the systemic setup, the plants were covered before spraying with a plastic tray leaving only the upper part/leaf tip (approximately 1 cm) of the first leaf uncovered. Using this approach we found that the CYP3RNA translocate from the first (sprayed) leaf into the second leaf as well as the shoot tissue over time (Fig. 4), the amount of sprayed dsRNA decreased from 1 d to 3 d after spray (das) application in leaves and shoots. Interestingly, the amount of dsRNA within the roots increased from 1das to 3 das (Fig. 5), indicating a translocation route from leaves to roots. However, to test whether the amount of transferred, SIGS-derived RNAs is sufficient to provoke target gene silencing, we inoculated the plants with *Fg* and measured the transcript level of *FgCYP51* target genes in leaves, shoots and roots of infected plants (Fig. 6). Notably, we measured the strongest target gene silencing in the first leaf, the leaf that was spray treated. However, the second leaf samples exhibited an overall target gene silencing of 80%, which is still very high. In the analysed shoot and root tissue we found 50% silencing of the fungal *CYP51* target genes. Together these data are consistent with our translocation observations, indicating that the transferred SIGS-associated RNAs can provoke silencing of their complementary target genes. Moreover, we found that 7 d after infection hypocotyls of plants sprayed with CYP3RNA developed less brownish lesions compared to TE-treated control plants suggesting that the amount of transferred RNA has the potential to prevent plants from *Fg* infection (Fig. 7).

**Figure 4:**
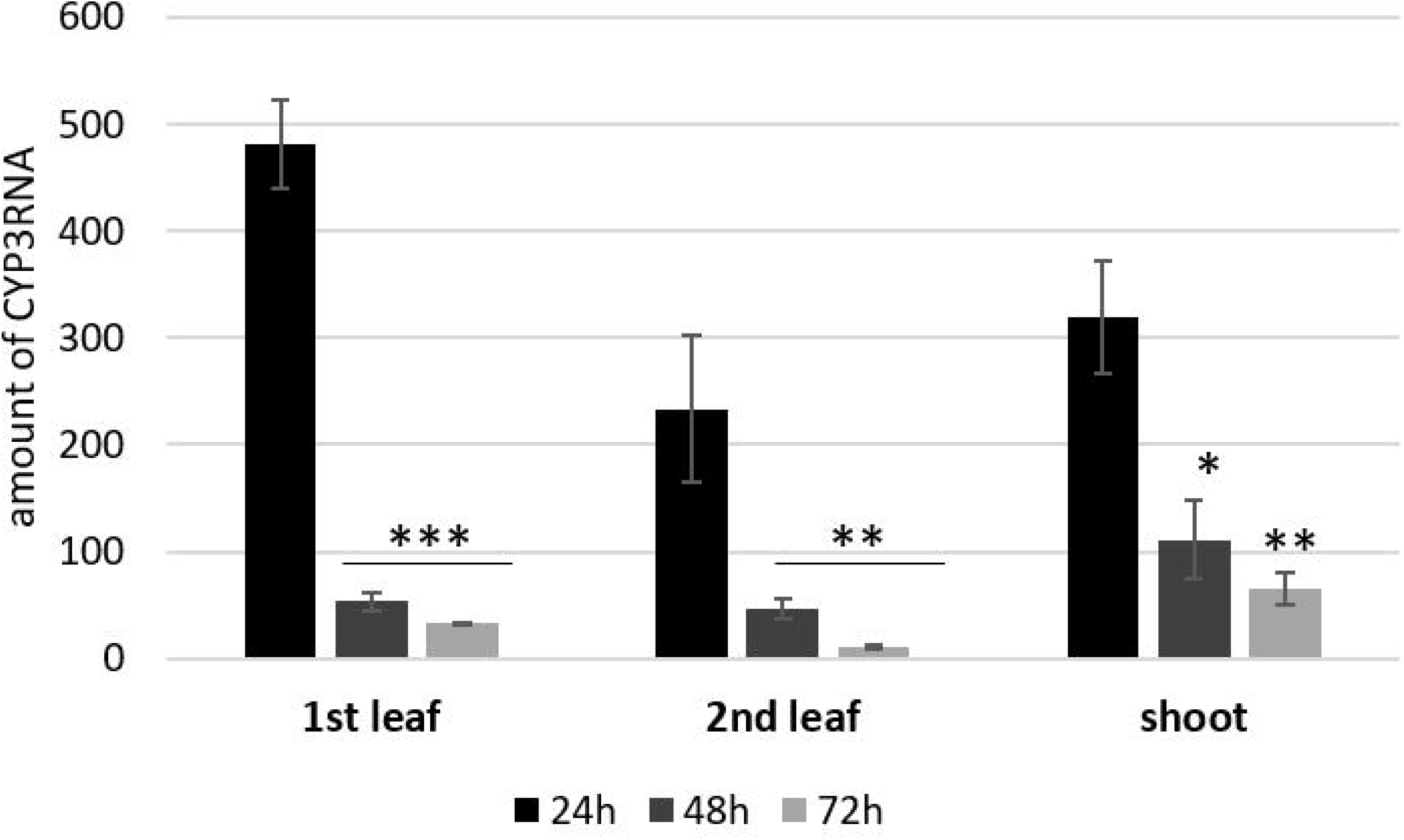
Detection of CYP3RNA in barley leaves and shoots at different time points after spray application. Amount of CYP3RNA in the first (sprayed) leaf as well as non-sprayed second leaves and shoot tissue. The relative amount of sprayed CYP3RNA measured by qPCR decreased over time in leaves and increased in roots compared to mock-sprayed control leaves. Bars represent mean values ± SEs of three independent experiments. The reduction of CYP3RNA vs. mock-sprayed leaves was statistically significant (*P < 0.05; Student’s t-test).

**Figure 5:**
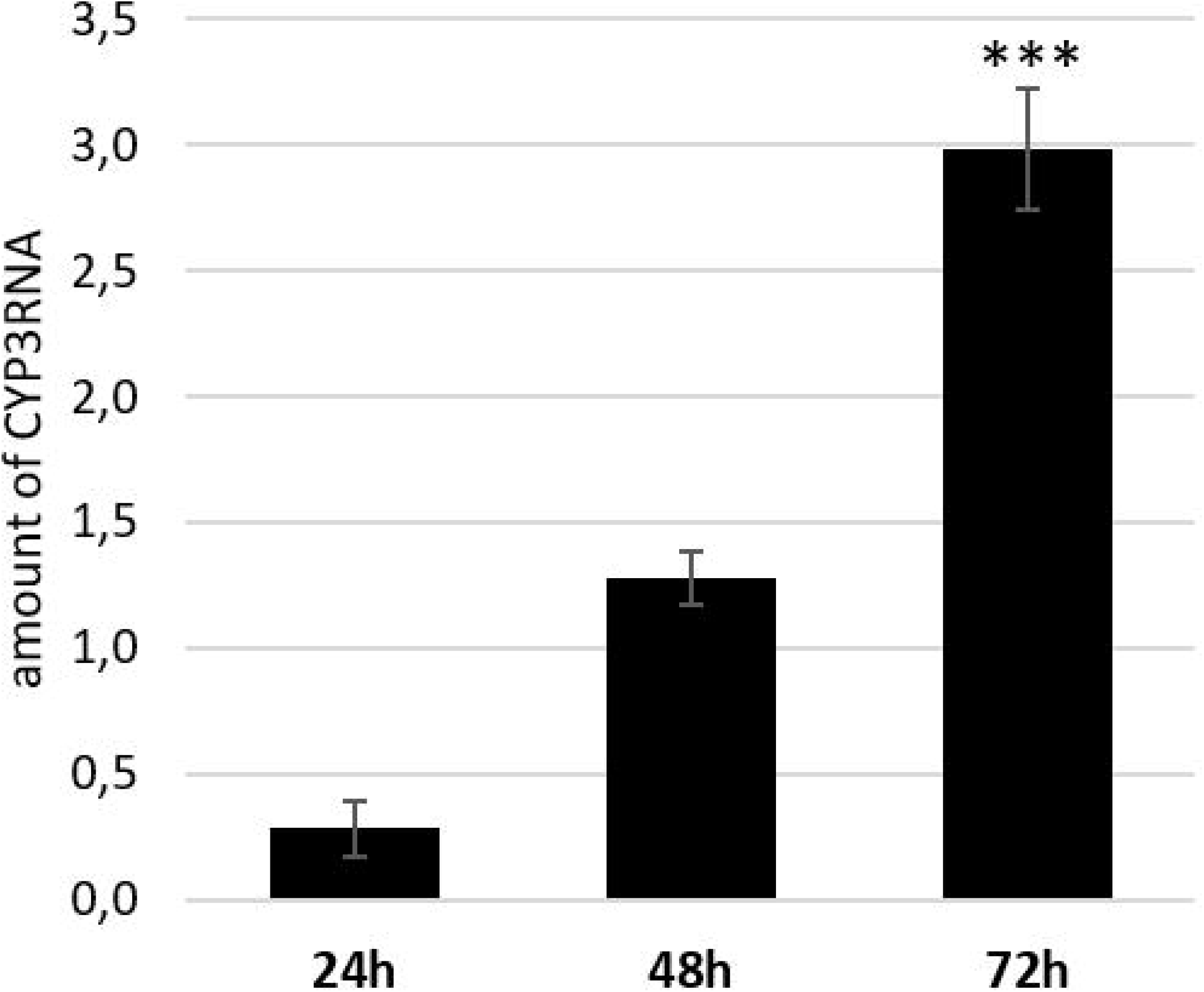
Detection of CYP3RNA in barley roots at different time points after spray application. Amount of CYP3RNA in the roots of barley plants 24h, 48h and 72h after spray treatment, respectively. The relative amount of sprayed CYP3RNA measured by qPCR decreased over time in leaves and increased in roots compared to mock-sprayed control leaves. Bars represent mean values ± SEs of three independent experiments. The reduction of CYP3RNA vs. mock-sprayed leaves was statistically significant (*P < 0.05; Student’s t-test).

**Figure 6:**
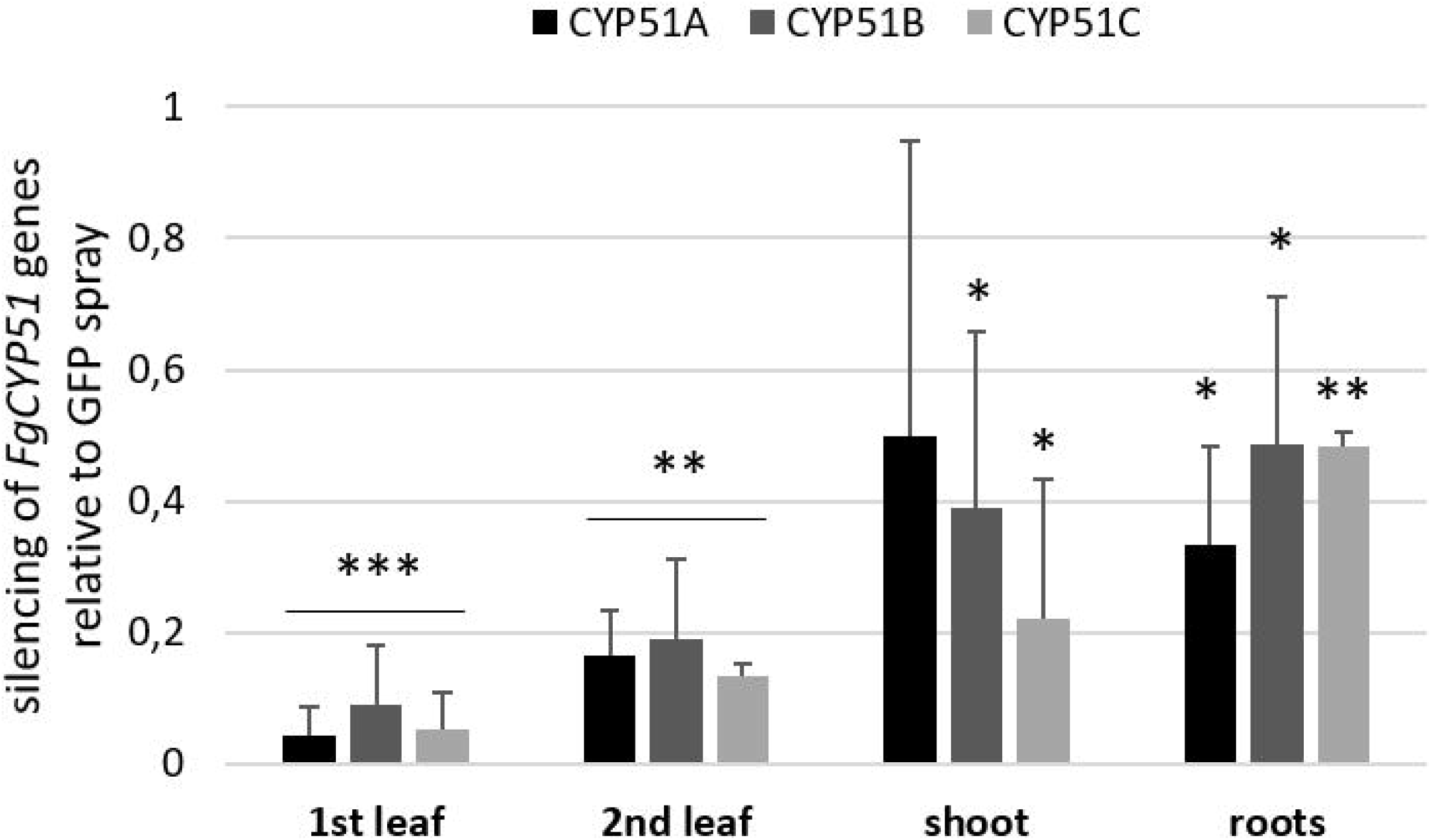
Gene-specific qPCR analysis of fungal *CYP51A, CYP51B*, and *CYP51C* transcripts at 6 dpi (corresponding to 9 d after spraying). The reduction in fungal *CYP51* gene expression on CYP3-dsRNA-sprayed leaves as compared with GFP-dsRNA-sprayed controls was statistically significant (*P < 0.05, **P < 0.01, ***P < 0.001; Student’s t test).

**Figure 7:**
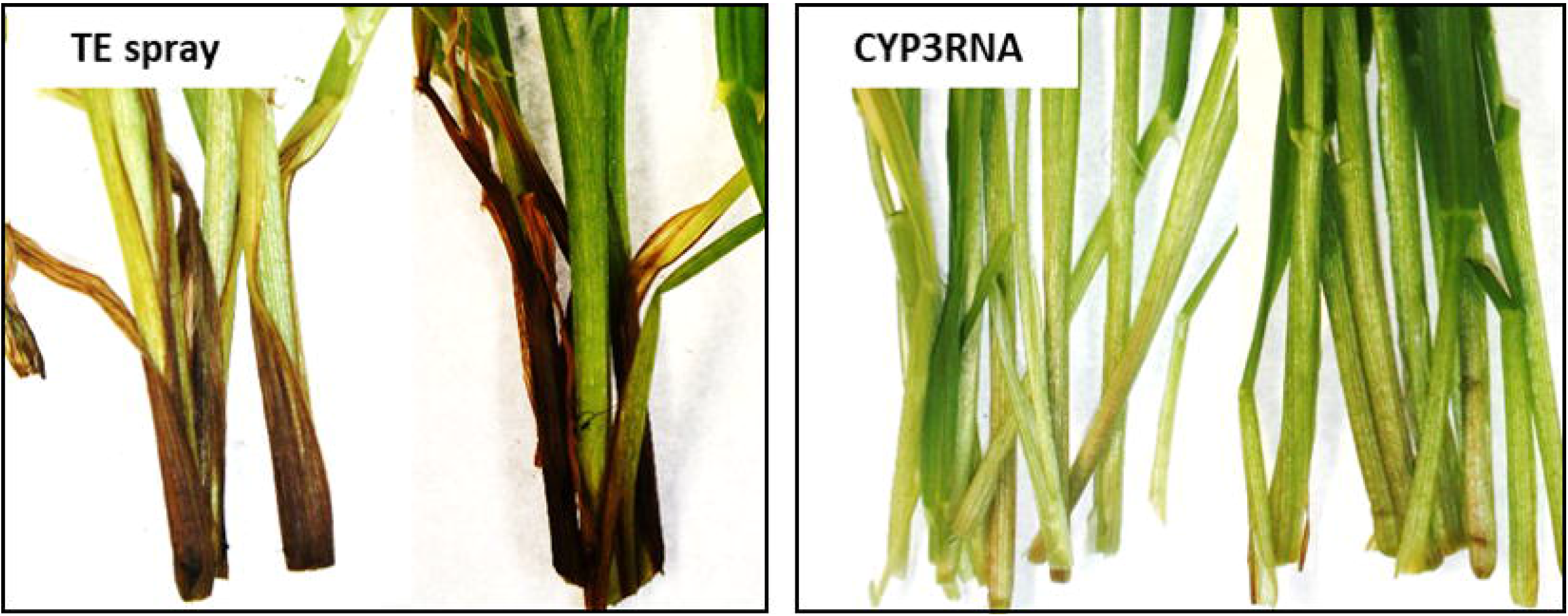
SIGS-mediated control of *Fg* on barley plants sprayed with CYP3RNA. (A) Two-week-old barley plants cv. Golden Promise were sprayed with CYP3RNA (20 ng μL^−1^) or TE-buffer, respectively. 72 hours later plants were inoculated with 2 × 10^4^ conidia mL^−^ of *Fg*. 7 d after infection hypocotyls of plants sprayed with CYP3RNA developed less brownish lesions compared to TE-treated control plants.

## Discussion

RNA sprays may provide an alternative strategy to avoid chemical pesticides and genetically modified crops for combating agricultural pests. However, systemic translocation of sprayed RNA biopesticides is critical for its successful future application in the field. To prove phloem-mediated transport of SIGS-associated RNAs we decided to transfer a HIGS approach established for a phloem-sap sucking grain aphid *Sitobion avenae* (Abdellatef et al., 2015) to a SIGS proof-of-concept study. To investigate uptake and transport of sprayed dsRNA, we tested whether locally sprayed Shp-dsRNA confers gene silencing in *S. avenae* infecting distal, non-sprayed segments of barley leaves. To this end, we assessed whether the phloem-transported inhibitory RNA affect *Shp* target gene expression in aphids fed on the dsRNA-sprayed barley leaves using qRT-PCR. The relative expression level of the aphid’s *Shp* gene was reduced by almost 60% compared to aphids that fed on buffer-sprayed controls (Fig. 1), suggesting that the transfer of inhibitory RNA from the plant phloem sap to the insect was successful. Next, we conducted another experiment to investigate whether the spray-applied Shp-dsRNA is translocated in the phloem and/or processed by the plant’s silencing machinery. Specifically, we used aphid stylectomy, a widely used technique to study physiological, mechanical and molecular properties of the plant phloem (Turgeon and Wolf 2009; Peel 1975; Thompson and Van Bel 2012). Here, we used this technique to gain access to the phloem sap of barley leaves (Fig. 2). To further explore the phloem-mediated transfer of sprayed Shp-dsRNA, we sprayed fluorescent labelled Shp-dsRNA_A488_ onto barley leaves using a systemic experimental design (Koch et al., 2016) following phloem sampling by stylectomy at the distal, non-sprayed leaf parts. Using CLSM, we detected a green fluorescent signal after cutting off the stylet tip of feeding aphids (Fig. 2 D). These results are consistent with our previous detection of the unprocessed 791 nt precursor CYP3RNA in both local and distal tissue using northern blot analysis, showing that the long dsRNA is systemically translocated within the plant (Koch et al., 2016). Moreover, investigation of longitudinal leaf sections revealed that the fluorescence was not confined to the apoplast but also was present in the symplast of phloem parenchyma cells, companion cells, and mesophyll cells, as well as in trichomes and stomata (Koch et al., 2016). Supportively, apoplastic movement of RNA has been proposed, e.g. to explain how maternally expressed siRNAs could be transferred from the endosperm of developing seeds into the symplastically isolated embryo (Martienssen 2010). However, the mechanism by which the sprayed RNA overcomes the apoplastic-sypmastic barrier is yet unknown.

Despite of the translocation of dsRNA, we found that also CYP3RNA-derived 21 nt and 22 nt siRNAs accumulated in the distal leaf segments, demonstrating that CYP3RNA was partly processed by the plant (Koch et al., 2016). Therefore, we predict that SIGS-derived siRNAs would also translocate via the barley phloem. To test this possibility, we additionally profiled Shp*-*dsRNA-derived siRNAs using stylectomy. Small RNA-sequencing analysis revealed Shp-dsRNA-derived siRNA in distal (non-sprayed) leaf segments (Fig. 3). These data suggest that Shp-dsRNA-derived siRNAs also are processed by the plant’s silencing machinery and are systemically transferred via the phloem. Importantly, we have previously shown that, when CYP3RNA_A488_-sprayed leaves were inoculated with *Fg* the fluorescent signal was also detectable inside fungal conidia, germtubes, and fungal mycelium (Koch et al., 2016). In addition or as an alternative to plant-mediated gene silencing in the fungus, systemic translocation within the plant and accumulation by the fungus could also mediate direct processing of CYP3RNA by the fungus’ gene silencing machinery to target fungal *CYP51* genes. This finding that unprocessed long dsRNA is absorbed from leaf tissue has important implications for future disease control strategies based on dsRNA. It is very likely that application of longer dsRNAs might be more efficient than application of siRNAs given their more efficient translocation. Moreover, in contrast to using only one specific siRNA, processing of long dsRNA into many different inhibitory siRNAs by the fungus may reduce the chance of pathogen resistance under field test conditions.

Consistent with our findings, there are several studies showing that mobile cell non-autonomous inhibitory RNAs that spread gene silencing into adjacent cells and tissues move through the vascular system (Lewsey et al., 2016, Palauqui et al., 1997). Grafting experiments revealed that siRNAs are detectable in tissues of DCL mutants that are defective for siRNA biogenesis (Molnar et al., 2010). This led to the speculation that siRNAs and not their long dsRNA precursors are the mobile silencing signals (Dunoyer et al., 2013). However, until now the exact molecular forms of mobile RNAs remain unclear (Parent et al., 2012). Here we showed movement of sprayed dsRNA from barley leaves over stems to the root tissue within three days after spray treatment (Fig. 4). Moreover, we found that the transferred SIGS-associated RNAs confer target gene silencing in the respective tissues as well as *Fg* disease resistance (Fig. 7, Koch et al., 2019). Interestingly, we measured the highest silencing efficiency in the tissue that was either directly sprayed or near to the sprayed locus. In other words, it seemed that the amount and/or the nature of the mobile RNA determine the systemic silencing efficiency. Therefore, we predict a dilution effect that correlate with the distance to the initial spray site (or tissue). This is consistent with another study, where the authors confirmed the spreading of dsRNA from local to systemic tissue by one hour after rub-inoculation of dsRNA using semi-quantitative RT-PCR. Moreover, they showed that dsRNA levels continuously decreased in the local (treated) tissue from 3 dpi to 9 dpi where dsRNA was no longer detectable (Mitter et al., 2017). More recently, Kaldis et al. (2018) showed that exogenously applied dsRNA derived from the silencing suppressor HC-Pro and the coat protein genes of zucchini yellow mosaic virus (ZYMV) protect watermelon and cucumber against ZYMV and spread systemically over long distances in cucurbits (Kaldis et al., 2018).

In summary, our data suggest that sprayed dsRNAs are taken up by the plant, spread systemically via the plant vascular system, and are partially processed into siRNAs by the plant’s gene silencing machinery (Fig. 8). As our results strongly support the notion of phloem-mediated long-distance movement of SIGS-associated dsRNA and/or siRNA, further research is needed to address central questions as: How are dsRNA transported at the apoplast-symplast interface? How does the fungus take up SIGS-associated RNAs from the apoplast and/or the symplast?

**Figure 8:**
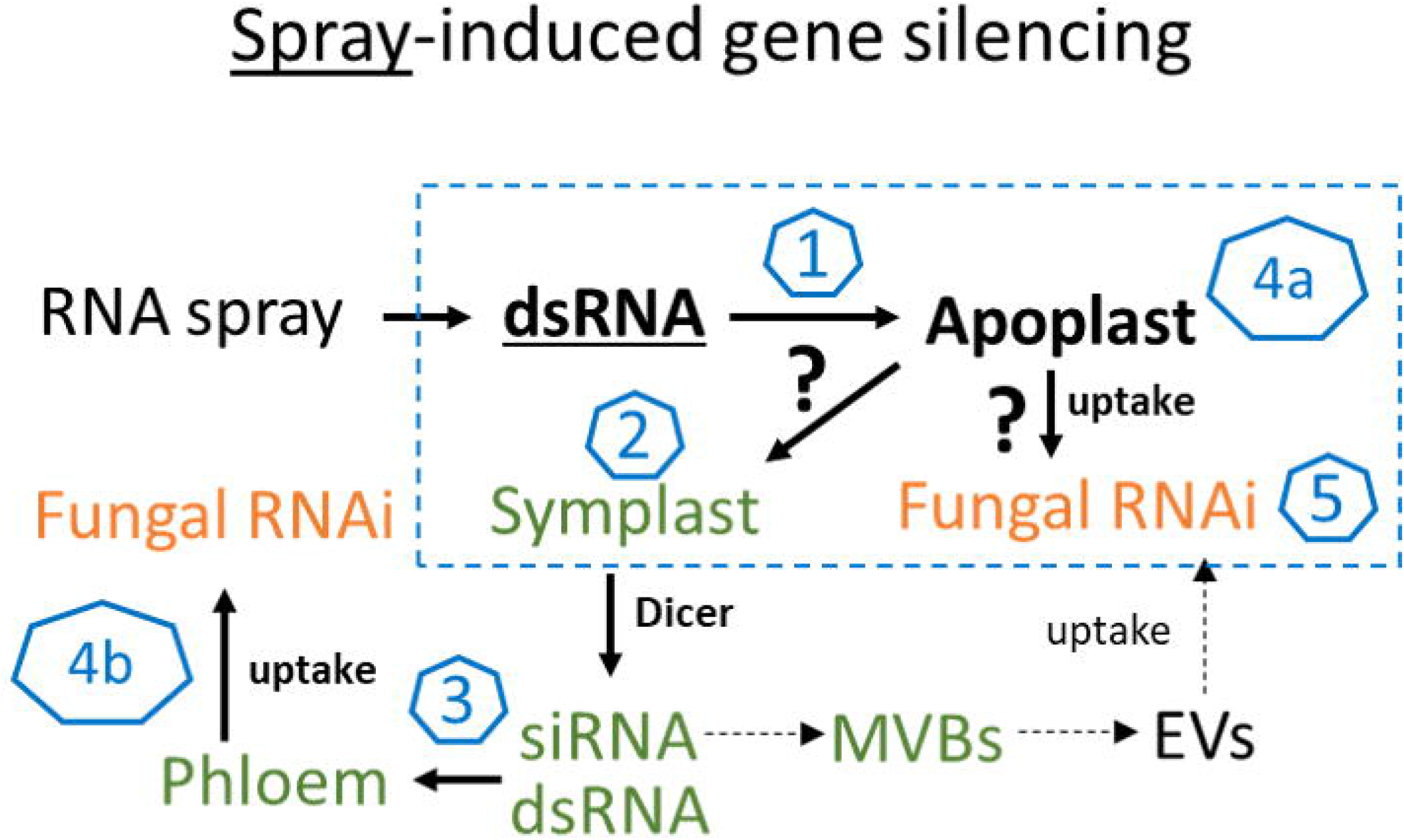
The molecular mechanism of SIGS is controlled by the fungal silencing machinery. In summary, our findings support the model that SIGS involves: (1) uptake of sprayed dsRNA by the plant (via stomata); (2) transfer of apoplastic dsRNAs into the symplast (DCL processing into siRNAs); (3) systemic translocation of siRNA or unprocessed dsRNA via the vascular system (phloem/xylem); (4) uptake of apoplastic dsRNA (a) or symplastic dsRNA/siRNA by the fungus (b); (5) processing into siRNA by fungal DCLs (Koch et al., 2016, Gaffar et al., 2019).

## Acknowledgment

We thank C. Birkenstock, U. Schnepp and V. Weisel for excellent plant cultivation. We also thank Dr. Jens Steinbrenner for assisting in microscopic analyses. This work was supported by the Deutsche Forschungsgemeinschaft, Research Training Group (RTG) 2355 (project number 325443116) to AK.

